# Awake Canine fMRI Predicts Dogs’ Preference for Praise Versus Food

**DOI:** 10.1101/062703

**Authors:** Peter F. Cook, Ashley Prichard, Mark Spivak, Gregory S. Berns

## Abstract

Dogs are hypersocial with humans, and their integration into human social ecology makes dogs a unique model for studying cross-species social bonding. However, the proximal neural mechanisms driving dog-human social interaction are unknown. We used fMRI in 15 awake dogs to probe the neural basis for their preferences for social interaction and food reward. In a first experiment, we used the ventral caudate as a measure of intrinsic reward value and compared activation to conditioned stimuli that predicted food, praise, or nothing. Relative to the control stimulus, the caudate was significantly more active to the reward-predicting stimuli and showed roughly equal or greater activation to praise versus food in 13 of 15 dogs. To confirm that these differences were driven by the intrinsic value of social praise, we performed a second imaging experiment in which the praise was withheld on a subset of trials. The difference in caudate activation to the receipt of praise, relative to its withholding, was strongly correlated with the differential activation to the conditioned stimuli in the first experiment. In a third experiment, we performed an out-of-scanner choice task in which the dog repeatedly selected food or owner in a Y-maze. The relative caudate activation to food-and praise-predicting stimuli in Experiment 1 was a strong predictor of each dog’s sequence of choices in the Y-maze. Analogous to similar neuroimaging studies of individual differences in human social reward, our findings demonstrate a neural mechanism for preference in domestic dogs that is stable within, but variable between, individuals. Moreover, the individual differences in the caudate responses indicate the potentially higher value of social than food reward for some dogs and may help to explain the apparent efficacy of social interaction in dog training.

## INTRODUCTION

As the first domesticated species, dogs have a unique relationship with humans. Dogs have been integrated into modern social life in many cultures, with millions serving as companion animals. As such, dogs benefit from a clear tendency of humans to bond socially with dogs (Beck & Madresh, 2008; Nagasawa et al., 2009; Odendaal & Meintjes, 2003; Stoeckel et al., 2014). But what is the nature of the relationship from the dog’s perspective? And, given the high degree of individual variability in dogs (see Scott & Fuller, 2012), how consistent across individuals are the biological underpinnings of this relationship? A better understanding of the proximal mechanisms driving dog-human interaction and the extent to which these vary across individuals will illuminate the dog-human social relationship. It is worth highlighting just how unique this cross-species relationship is. While commensalism and symbiosis are not uncommon in the animal kingdom, a species-wide extension of social bonding mechanisms to include a wholly unrelated species is apparently very rare, and raises the intriguing possibility that human social behavior has served as a strong adaptive pressure in the evolution of domestic dog sociobiology (Reid, 2009). Quantifying the relative value of food versus praise would also help inform ongoing and contentious debates regarding the most effective methods in dog training (e.g., Blackwell et al., 2008; Hiby et al., 2004; McKinley & Young, 2003)

Dogs are gifted at attending to, and interpreting, subtle human social cues (Lakatos et al., 2012; Merola, Prato-Previde & Marshall-Pescini, 2012; Muller et al., 2015), and a behavioral literature suggests that dogs act as if socially attached to humans (Topal et al., 1998; Palmer & Custance, 2008; although see: Prato-Previde et al., 2003; Rehn, McGowan & Keeling, 2013). Despite this, the motivations behind dog behavior toward humans can be difficult to disentangle from behavior alone. In terms of measuring preference, dog social behaviors are highly susceptible to prior patterns of food reinforcement (Bentosela et al., 2008; Elgier et al. 2009), and dogs frequently treat interaction with their owner as an avenue to acquire food (Cook, Arter & Jacobs, 2014), even suppressing interest in food under communicative situations (Pongracz et al., 2013). In direct tests of behavioral preference, some dogs select their owners and others food (Gacsi et al., 2005; Topal et al., 2005; Feuerbacher & Wynne, 2014; Feuerbacher & Wynne, 2015) – but the behavior appears to be contingent on testing method, socialization history, reinforcement history and potentially many other factors including attention, stimulus salience, and satiety. Further, while social reinforcement is a commonly used tool in dog training (Hiby, Rooney & Bradshaw, 2004), and many trainers believe it to be effective, it is quite difficult to experimentally isolate social and food reward in a training paradigm to measure their relative contribution to learning. Food delivery in dog training almost always includes a social component, and the acquisition rate of new behaviors can vary greatly depending on factors aside from reinforcement type. Notably, animals have long been known to show faculty for generalized learning (see Harlow, 1949), meaning that novel behaviors may be learned more quickly due to prior experience with even tangentially related learning tasks. The difficulties of isolating variables contributing to choice behavior and learning rate in purely behavioral paradigms highlights the potential value of a neurobiological approach seeking a consistent signal underlying individual differences in behavior.

Recent findings indicate that oxytocin, a neuropeptide critical for pair bonding within some species (Winslow, 1993; Young & Wang, 2004) has a role in mediating dog behavior toward humans (Odendaal & Meintjes, 2003; Romero et al., 2014; Nagasawa et al., 2015; Thelke & Udell, 2015). While oxytocin supports accounts of social attachment (although see Walum et al., 2016), the proximal neural mechanisms of social reward in the dog are unknown. In humans and other mammals, oxytocin receptors are dense in the ventral striatum, including the nucleus accumbens and caudate nucleus (Fruend-Mercier et al, 1987; Olazabal & Young, 2006; Ross & Young, 2006), and these brain regions are also involved in social attachment in humans and other species (Rilling et al., 2002; Izuma, Saito & Sadato, 2008; Young, Liu, & Wang, 2008; Burkett et al., 2011). There is also strong evidence of a dissociation between the ventral and dorsal striatum, such that the ventral portion is more relevant to reward anticipation and learning (O’Doherty et al., 2004). In humans, activity in the ventral striatum has been associated with a wide variety of rewards, including both monetary and social (Delgado, 2007; Haber & Knutson, 2010; Lin, Adolphs & Rangel, 2012; Pauli et al., 2016), and activation increases with increasing valuation (Knutson et al., 2000; Koeneke et al., 2008; Howe et al., 2013). Ventral striatal activity has also been shown to predict purchases (Knutson & Bossaerts, 2007), the choice of cola-drinks (McClure et al., 2004), and the popularity of songs (Berns et al., 2012). In animals, the ventral striatum has been shown to code the relative value of competing outcomes (Cromwell, Hassani & Schultz, 2005). Thus, there is good reason to believe that the analogous structure in dogs could be used to measure stable individual differences in relative preference for food and social rewards.

The advent of awake canine neuroimaging (Berns, Brooks & Spivak, 2012; Andics et al., 2014; for review see Cook et al., 2015) provides a unique opportunity to probe the neural mechanisms underlying the dog-human bond. Because of its coding of reward value, ventral caudate activation can serve as a measure of “choiceless utility” (Loomes & Sugden, 1982), that is, the value a stimulus has for an individual who is not acting to acquire it. Choiceless utility tasks control for many of the confounds influencing active choice in dogs, and thus may be optimal for assessing the extent to which dogs value social interaction.

In dogs, we have previously shown a temperament-dependent increase in neural activity in the ventral caudate when dogs are presented with a stimulus associated with incipient receipt of food reward (Cook, Spivak & Berns, 2015) and when they are presented directly with olfactory stimuli associated with familiar humans without linked reward (Berns, Brooks & Spivak, 2015). While other brain regions may be differentially activated by reward preferences, no other region has the same strength of *a priori* justification for use with dogs (or any other animal) as does the ventral striatum (c.f. Ariely & Berns, 2010 for Bayes’ factor of reward and striatum). If dogs are socially attached to humans and value interactions with them for more than the provision of food, social reward should be represented in ventral caudate activation in expectation of, and response to, human interaction. Moreover, the strength of this activation, relative to food, could be used as a marker for social preference. If so, then ventral caudate activation should be predictive of behavioral choice between food and social interaction.

In the present study, we conducted three independent experiments. In Experiment 1, to identify a stable neural response associated with relative valuation of social interaction and food, we used fMRI to measure ventral striatal activation in 15 awake, unrestrained dogs while they passively viewed objects associated with incipient food reward or receipt of verbal praise from a primary handler. A neutral object, not previously associated with reward outcome, served as a control condition. Consistent with a choiceless utility approach, the primary experimental comparison was between activation during presentation of the food-predicting and praise-predicting conditioned stimuli, prior to delivery of reward.

In Experiment 2, to determine stability of the neural valuation of praise within individual dogs, we replicated Experiment 1 with one alteration: on a subset of praise trials, the owner did not appear and praise the dog after the praise-predicting object was displayed. This withholding of praise constituted a violation of expectation, or a “negative prediction error,” as previously studied in humans (Schultz & Dickinson, 2000; McClure, Berns & Montague, 2003). Negative prediction errors result in a decrement of caudate activation, the magnitude of which is typically understood to relate to the value of whatever is being withheld. Experiment 2 allowed us to validate the relative neural response to praise and food from Experiment 1 using a complimentary experimental approach and a different temporal component of reward processing (expectation violation). Dogs who more greatly valued social reward ought to show greater positive activation to an object predicting incipient social reward (Experiment 1) and greater positive activation to receipt of praise versus withholding of expected praise (Experiment 2).

In Experiment 3, to determine whether any stable neural marker of valuation could explain behavioral variability in active choice, each of the subjects took part in an independent, out-of-scanner behavioral choice task where they repeatedly selected between receipt of food or social interaction with their primary handler. Responses were modeled as relative stay and shift probabilities between food and handler, which could then be directly compared to relative caudate activations from Experiment 1.

## METHODS & MATERIALS

### Participants

Participants for Experiments 1 and 3 were 15 domestic dogs, and all dogs/owners were volunteers from the Atlanta area. Thirteen of these dogs completed Experiment 2. All dogs had successfully completed at least one prior brain scan and had received extensive training to hold still in the scanner (see Berns, Brooks & Spivak, 2012).

### Training

Because participants were already skilled MRI dogs, the only additional training for the current study was to condition association of outcome with the three object stimuli used in both in-scanner experiments. The three objects were a toy car, a toy horse, and a hair brush, and they were associated with verbal praise, food reward, and nothing respectively. Training for each subject was conducted in two sessions on separate days at our training facility within two weeks of live scanning for Experiment 1. During training sessions, dogs were stationed in their custom-made chin rests inside our mock MRI coil. An out-of-sight experimenter visually presented the three objects on the end of long sticks approximately 2 feet from the dog’s nose. Each object was presented for 10 seconds and was always followed by its associated outcome (Fig. 1, Movie 1). After the car had been presented, the dog’s primary handler came into view and offered 3 seconds of verbal praise before exiting the dog’s field of view. After the horse had been presented, a hot dog piece was provided to the dog on the end of a feeding stick, with no human coming into view. After the brush had been presented, nothing occurred. Inter-stimulus intervals were approximately 5-10 seconds. A training session was 60 trials, such that each object and its outcome were presented 40 times during training across the two sessions. The order of presentation was randomized with the constraint that no object-outcome pair was presented more than three times in a row.

**Figure 1:**
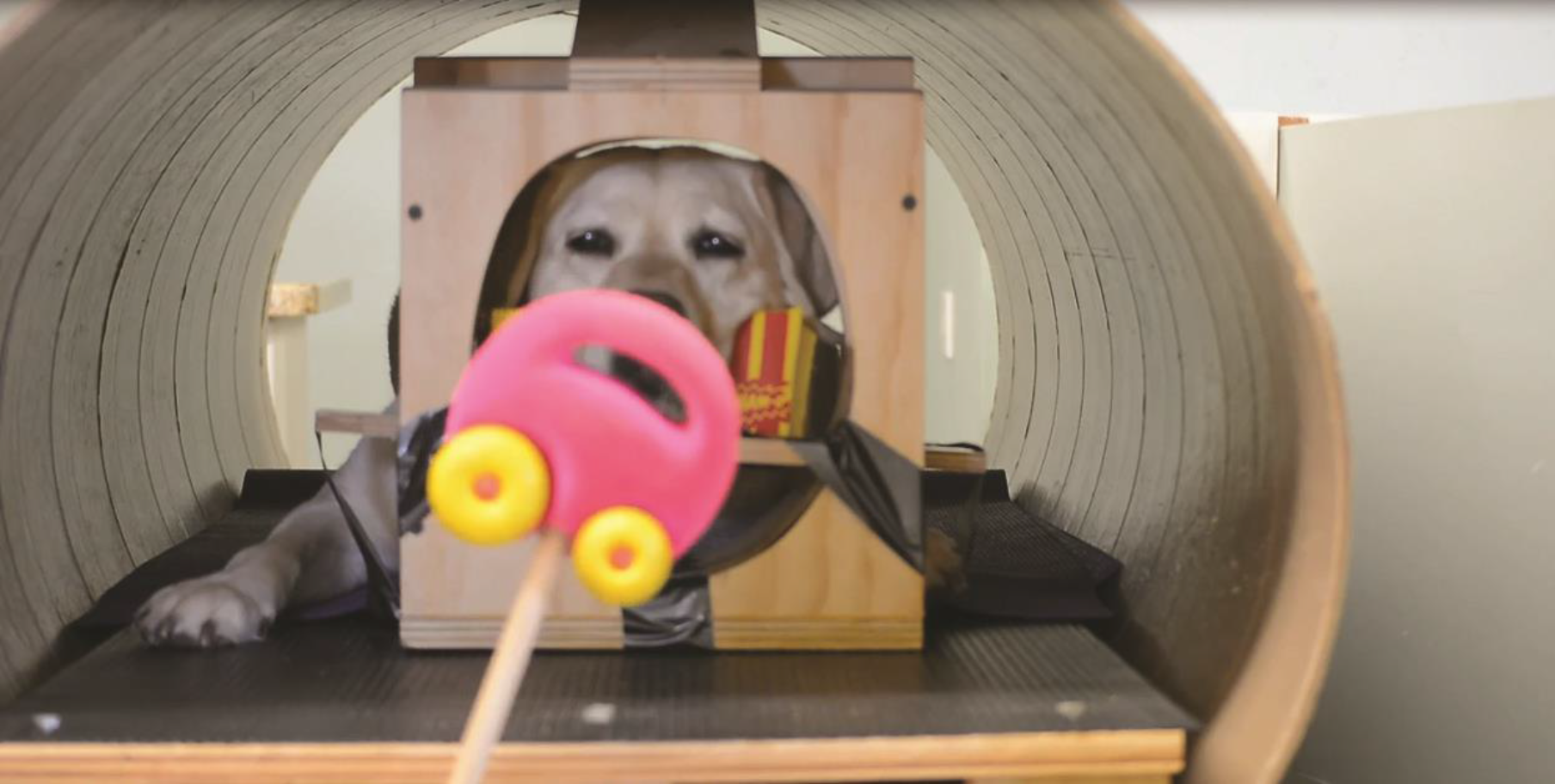
In-scanner reward expectation task. Subjects were presented with the three experimental stimuli in the mock scanner during training. Subject Kady is shown here viewing the car object during conditioning (see also Movie 1). Each dog was exposed to each stimulus and its outcome (car-praise, horse-food, and brush-nothing) 40 times in a pre-set semi-random schedule during training within the two weeks prior to live scanning. They were exposed to each object and paired outcome 8 more times immediately prior to live-scanning. During live scanning, objects were presented for 10 seconds each, followed immediately by outcome.

### Imaging

All basic procedures and imaging parameters were identical between the two separate functional MRI experiments. Scans were conducted during the late morning or early afternoon, and owners were instructed not to give their dogs a large meal immediately prior to scanning. All dogs would have eaten breakfast. Prior to both Experiment 1 and Experiment 2, each dog did a “dry run” of the scanning procedure, stationing in their custom chin rest in the scanner bore and receiving each object-outcome pairing 8 times in a pre-set random schedule. This was to insure that the dogs remembered the pairings before live scanning.

As in our prior experiments (see Berns, Brooks & Spivak, 2013), during live-scanning, all participants wore ear protection, either Mutt Muffs^TM^ or ear plugs with vet wrap, depending on dog and owner preference.

All scans were acquired on a 3T Siemens Trio MRI, and scan parameters were similar to those in Cook, Spivak & Berns (2014). Functional scans used a single-shot echo-planar imaging (EPI) sequence to acquire volumes of 23 sequential 2.5 mm slices with a 20% gap (TE=25 ms, TR = 1200 ms, flip angle = 70 degrees, 64×64 matrix, 3 mm in-plane voxel size, FOV=192 mm). A T2-weighted structural image was previously acquired during one of our earlier experiments using a turbo spin-echo sequence (25-30 2 mm slices, T = 3940 ms, TE = 8.9 ms, flip angle = 131 degrees, 26 echo trains, 128x128 matrix, FOV=192 mm). 1500 to 2000 volumes were acquired for each subject in both Experiments 1 and 2, equaling 30-40 minutes of scan time for each procedure.

### Experimental Design

Stimulus presentation method and event recording were identical in Experiments 1 and 2. Stimuli were presented as during training, on the end of sticks by an out-of-view experimenter. Food (pieces of hot dog) was provided after horse trials on the end of a long stick with no human in view. The dogs’ primary handlers were stationed just out of site to the left of the magnet bore from the dog’s perspective, and appeared in order to praise the dog after presentation of the car object.

Trial events (onset and offset of object presentations) were recorded by an observer out-of-sight of the subject via a four-button MRI-compatible button-box. A computer running PsychoPy (Peirce, 2009) was connected to the button-box via usb port, and recorded both the button-box responses by the observer and scanner sequence pulses.

#### Experiment 1

Each of the 15 subjects received 32 presentations of each stimulus-outcome pairing (car-praise, horse-food, brush-nothing) in a randomized schedule constrained such that no stimulus could be presented more than three times in a row. Depending on comfort in-scanner, subjects were scanned in 2, 3, or 4 separate runs on the same day.

#### Experiment 2

Each of the 13 subjects were scanned within two months of completing Experiment 1, with no extra training related to this study. One dog (Ozzie) moved too much during scanning, polluting the fMRI data, so his results were not used in subsequent analyses. In live-scanning, the other 12 subjects received between 12 and 16 presentations of the horse-food pairing, 15 and 20 presentations of the brush pairing, and between 60 and 80 presentations of the car. On 1/4^th^ of the car presentations, the expectation violation trials, the owner did not appear and praise the dog. The range in trial number was due to two of the 12 dogs who began to shift positions in the scanner and so completed only 3/4ths of the planned trials.

### Functional Data Preprocessing and Analysis

Image preprocessing and analysis—including motion correction, censoring, normalization, and GLM fitting—used AFNI (NIH, Cox 1996) and its affiliated tools, and was carried out for both Experiments 1 and 2 as in our previous studies (e.g., Cook, Spivak & Berns, 2014). Briefly, 2-pass, 6-parameter affine motion correction was used with a hand-selected reference volume for each dog. Because of excessive motion, one dog had to have one run of 24 trials discarded in Experiment 1. Dogs moved while consuming food (but not typically praise), and in-between trials, so data were censored, relying on a combination of outlier voxels in terms of signal intensity (greater than 1% signal change from scan-to-scan) and estimated motion (greater than 1mm displacement scan-to-scan). Visual inspection of the censored volumes was conducted to be certain that all bad volumes (e.g., when the dog’s head was out of the scanner) had been excluded. In Experiment 1, on average, 54% of total EPI volumes were retained for each subject after censoring (ranging from 30% to 70%). Fifty-four percent of volumes were also retained on average for Experiment 2 (ranging from 40% to 72%). This was consistent with previous experiments with this cohort of dogs (Berns, Brooks & Spivak, 2013; Berns, Brooks & Spivak, 2014). EPI images were smoothed and normalized to %-signal change with 3dmerge using a 6 mm kernel at Full-Width Half-Maximum (FWHM).

In both Experiments 1 and 2, for each subject, motion-corrected, censored, smoothed images were put into a General Linear Model estimated for each voxel using AFNI’s 3dDeconvolve. In both GLMs, motion time courses generated by motion correction were included as regressors, as were constant and linear drift terms. We have previously measured hemodynamic response function (hrf) in dogs to peak between 4-6 seconds (Berns, Brooks & Spivak, 2012). This is similar to what is seen in the human literature, so our models incorporated typical hrfs with a single gamma function using default parameters in AFNI.

#### Experiment 1

Task related-regressors were: (1) presentation of the car (praise) object, (2) presentation of the horse (food) object, and (3) presentation of the brush (nothing) object. These three task regressors were modeled durationally using AFNI’s dmUBLOCK function.

#### Experiment 2

Task related regressors were: (1) onset of praise, (2) onset of violation of expected praise, (3) offset of the brush object, (4) presentation of the horse (food) and car (praise and no-praise) objects, and (5) presentation of the brush (nothing) object. Regressors 1-3 were modeled as impulse response functions, convolved with a single gamma function approximating the hrf. These regressors included both the onset of praise and the onset of violation of expected praise. In both cases, “onset” was coded as one second after the removal of the praise-predicting object (car) such that hemodynamic response was measured peaking over the period that praise was offered or immediately following violated expectation. Regressors 4 and 5 were modeled durationally with AFNI’s dmUBLOCK function.

### Region of Interest (ROI) Analysis

Because our interest was in reward value as coded in the ventral striatum, all quantitative analyses based on imaging results used activation values in the ventral striatum. While portions of the ventral striatum other than the caudate are relevant for reward (notably the nucleus accumbens), we cannot resolve a structure this small given the resolution of both the structural and functional scans and the size of the dog’s brain. As in Berns, Brooks & Spivak, 2014, regions of interest (ROIs) were drawn separately on the left and right ventral caudate on each subject’s structural scan (Fig. 2A & 2B). At resolution, we assume these ROIs also include the nucleus accumbens. These ROIs were defined *a priori* using anatomical criteria, avoiding confounds and multiple comparison problems common to whole-brain imaging analyses and group spatial normalization. Statistical maps were transformed into structural space at the individual level, and mean beta values for regressors-of-interest were extracted for each ROI, representing mean percent signal change for the related experimental conditions. To probe for a main effect of reward signals in Experiment 1, we computed a t test on the beta values from the contrast [((CS_praise_ + CS_food_)/2) – CS_neutral_]. We also compared beta values from Experiment 1 and 2 for [CS_praise_ – CS_neutral_] and [CS_food_ – CS_neutral_] in a linear mixed effects model. To establish consistency between valuation of praise in Experiment 1 and Experiment 2, we computed a linear regression on the beta values from the contrasts: Exp1[CS_praise_ – CS_food_] and Exp2[Praise – No Praise].

**Figure 2:**
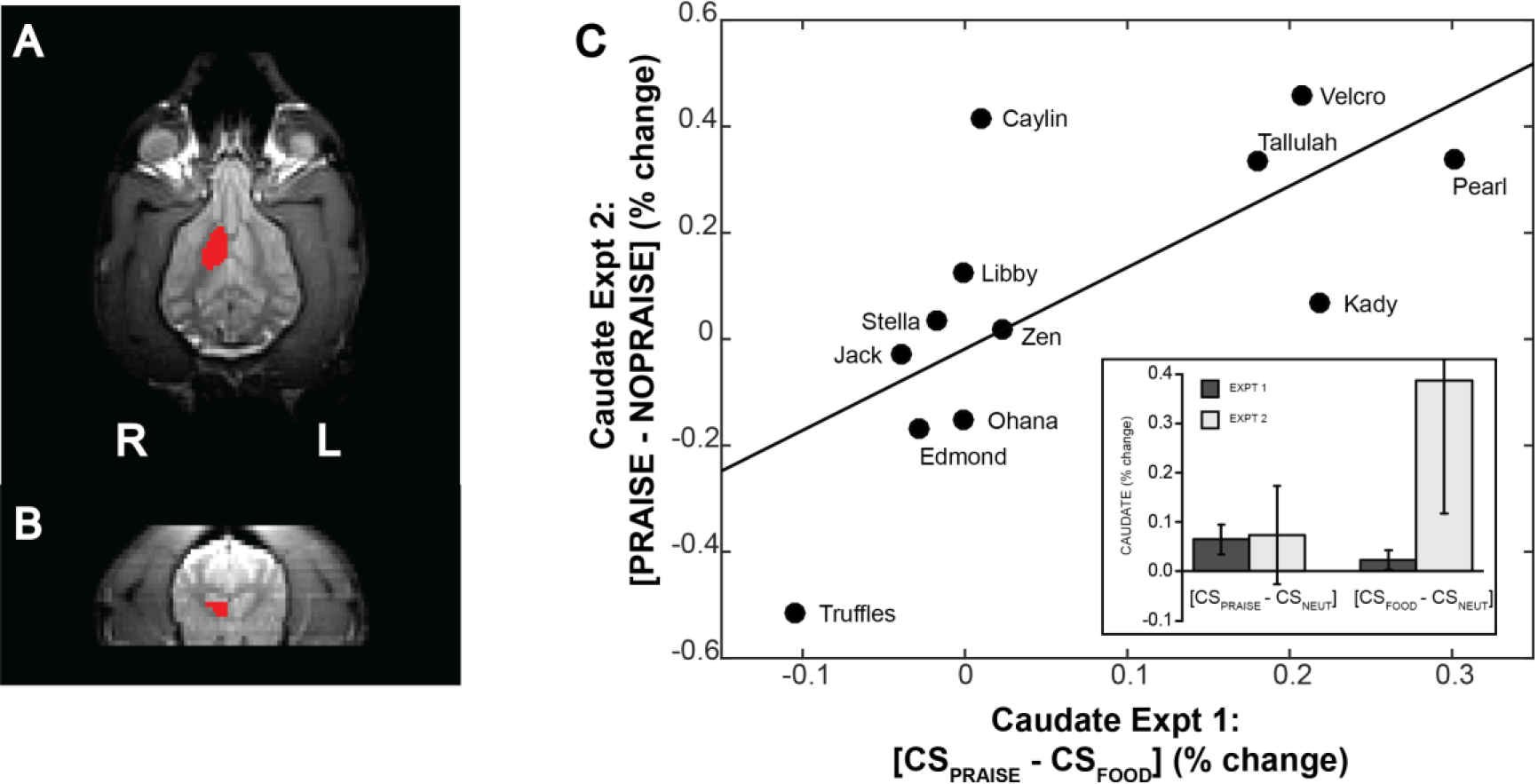
Structural definition of ventral caudate and demonstration of consistent neural valuation. (*Left*) For each subject, a mask of ventral left and right caudate nucleus was drawn directly on the T2 structural scans, shown here in the transverse (*A*) and coronal (*B*) planes. In the transverse plane, the rostral portion of the brain is toward the top of the image where the eyes are visible laterally and the olfactory bulb medially. Caudate masks were drawn to be consistent with those we have used in previous studies and covered the entirety of the head of the caudate nucleus ventral to the ventral-most portion of the genu of the corpus callosum. To obtain condition and contrast specific measures of BOLD activation in these regions, each individual’s statistical maps in functional space were transformed to structural space, and the relevant beta values were averaged across the caudate ROIs. (*C*) Regression line fitting differential caudate activation to expectation of praise versus food (Experiment 1, x axis) versus differential caudate activation during receipt of praise versus withholding of praise (Experiment 2, y axis). (F(1,10) = 9.49, R^2^ = 0.49, P = 0.01, two-sided). (*C – inset*) Mean percent signal change in the *a priori* ventral caudate masks is shown for the contrasts [CS_praise_ – CS_neutral_] and [CS_food_ – CS_neutral_] for Experiments 1 and 2. Relative response to food cue versus neutral cue and praise cue versus neutral cue were not significantly different within or between Experiments 1 and 2, nor was there an interaction between experiment and contrast (Experiment: F_1,14.6_ = 1.78, P = 0.20; Contrast: F_1,14.6_ = 0.95, P = 0.35; Experiment × Contrast: F_1,14.6_ = 1.61, P = 0.22).

### Experiment 3

#### Behavioral Testing

Each dog was tested in a familiar environment at our training facility. Dogs were tested in the afternoon, and, as with scanning, owners were instructed not to feed their dogs immediately prior to testing. Testing used a Y-maze setup, with the owner, seated with back to the entrance to the maze, and a bowl of food at the end of the arms, 12 meters from the entrance to the testing enclosure (Fig. 3, Movies 2 & 3). Tests began with four trials of warmup in which each dog was restricted to choosing owner, food, food, and owner in that order, balanced between the left and right arms of the Y maze. In between trials, the dog was held in a waiting room. There were then 20 data trials in which each dog was released into the testing enclosure and free to make her choice. When the dog approached the handler, the handler gave the dog verbal praise and petting. When the dog approached the food bowl, she was free to eat the contents, between one and three small pieces of Pupperoni (^®^Big Heart Pet Brands). Choices were coded in real time by a hidden observer. After a choice, or after 20 seconds of no choice, the dog was collected by an experimenter and returned to the waiting room. Approximate time in the room between trials was 15 seconds. At initial testing, two of the 15 subjects never selected the food reward, suggesting that they did not understand its continued availability. Both were re-tested on a later day with an extended warmup phase (10 trials balanced between forced choice of food and owner) and results from follow-up testing were used.

**Figure 3:**
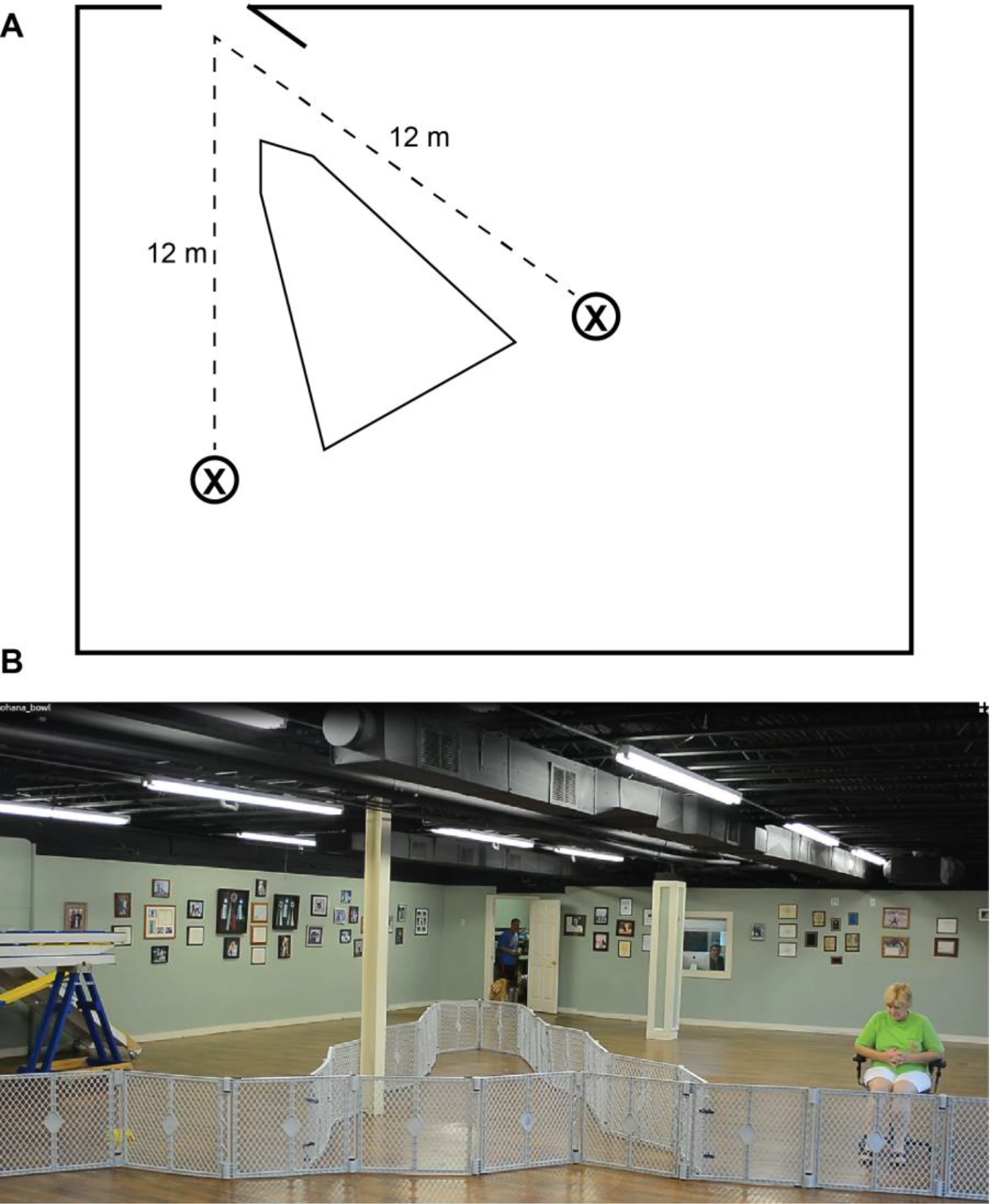
Behavioral preference test. (A) schematic representation of Y maze testing set up. Dogs were released from the control room door at a point equidistant from the owner and food locations (X). (B) Subject Ohana being released from the control room to make her choice between food (yellow bowl at left) and owner (at right) (also see Movies S2 and S3). On each trial, the dog was allowed to make one selection, resulting in either consuming the food or receiving verbal praise and petting from the owner. Location of owner and food was switched on each trial to control for effect of side biases. Following release from the control room, each dog was allowed 20 seconds to make one (and only one) choice before being collected and returned to the control room for another trial.

### Hidden Markov Model

Because preferences may shift during the course of a sequential choice task, and because some dogs didn’t make choices on all trials, simple proportions of responses may fail to capture the underlying dynamics of choice. We hypothesized that the dog’s choice was mediated by a “hidden” state within her brain; therefore, we used a simple, two-state, Hidden Markov Model (HMM, Rabiner, 1989; Eddy, 1996) to model sequential choice. We assumed that the dog was in a either “food” or “praise” state and could shift or stay between them. When in the “food” state, a behavior was emitted toward the bowl with probability, E1. When in the “praise” state, behavior was emitted to the owner with probability E2. State transition probabilities (P_PP_, P_PF_, P_FF_, P_FP_) and emission probabilities (E_1_, E_2_) were estimated in Matlab using *hmmtrain.* Initial transition probabilities were set to: [0.5 0.5; 0.5 0.5], and initial emission probabilities were set to 0.9 for food→bowl and praise→owner, and 0.1 to ‘nothing,’ which allowed for trials in which a dog did not make a choice. The difference between owner-stay and food-stay probabilities was then used as an index of behavioral preference.

### Comparison of Imaging and Behavior

To determine the stability of neural responses for food and praise, difference in mean caudate activation to the praise and food objects in Experiment 1 was compared to difference in mean caudate activation during receipt of praise and violation of praise expectation in Experiment 2 using linear regression. To determine whether the neural response could predict choice behavior, the difference in mean caudate activation to the praise and food objects in Experiment 1 was compared to the difference in transition probabilities between owner and food in the Experiment 3. To fit the logit model, we first transformed the difference in transition probabilities into an equivalent number of owner/bowl choices, according to: N(P_pp_-P_ff_+1)/2, where N was the total number of trials in which a dog made a choice. The logit model was then fit in R using a probit link function. We used logistic regression to account for the bounded nature of the transition probabilities.

## RESULTS

In Experiment 1, average activation to the food and owner objects was greater than activation to the neutral object (*i.e.,* defacto baseline) across all dogs [(CS_praise_ + CS_food_)/2) – CSneutral], indicating that the dogs had learned the reward associations with the test objects (T(14)=2.42, P=0.03, two-sided, Cohen’s d = 0.63). The ventral caudate was active to both CSs and showed roughly equal activation (i.e. within one s.d. of the group mean) or greater activation to praise versus food cues in 13 of 15 dogs. However, average ventral caudate activation did not differ between the owner and food conditions [CS_praise_ - CS_neutral_ vs CS_food_ – CS_neutral_] (T(14)=1.28, P=0.22, two-sided).

Relative response to food versus neutral and praise versus neutral were not statistically distinct within or between Experimenst 1 and 2, nor was there an interaction between experiment and contrast (Experiment: F_1,14.6_ = 1.78, P = 0.20; Contrast: F_1,14.6_ = 0.95, P = 0.35; Experiment × Contrast: F_1,14.6_ = 1.61, P = 0.22) (Fig. 2C).

As hypothesized, we found that differential caudate response to CSs predicting praise versus food (Experiment 1 [CS_praise_ – CS_food_]) was strongly predictive of differential caudate response to receipt versus withholding of praise (Experiment 2 [Praise – No Praise]) (F(11) = 9.49, R^2^ = 0.49, P = 0.01, two-sided) (Fig. 2C). Thus, in accordance with both Thorndike’s Law of Effect and the theory of choiceless utility, there was a strong relationship between the relative value of expected praise versus expected food (Experiment 1) and the value of received praise versus praise-withholding (Experiment 2). Measured across two independent experiments, and across reward expectation and reward receipt, these results suggest a consistent neural signal of valuation in the dogs’ ventral caudate.

Differential caudate activation to the owner versus food object in the passive in-scanner task [CS_praise_ – CS_food_] was also strongly correlated with the owner-stay probability relative to the food-stay probability in a logit model (Z(13) = 6.24, P < 0.001, two-sided, null deviance = 93.08, residual deviance = 50.10, chi square goodness of fit P < 0.001)(Fig. 4C). In other words, passive neural preference strongly predicted active choice, with the ventral caudate acting as the hidden variable that biased the dogs’ choices of food or praise.

**Figure 4:**
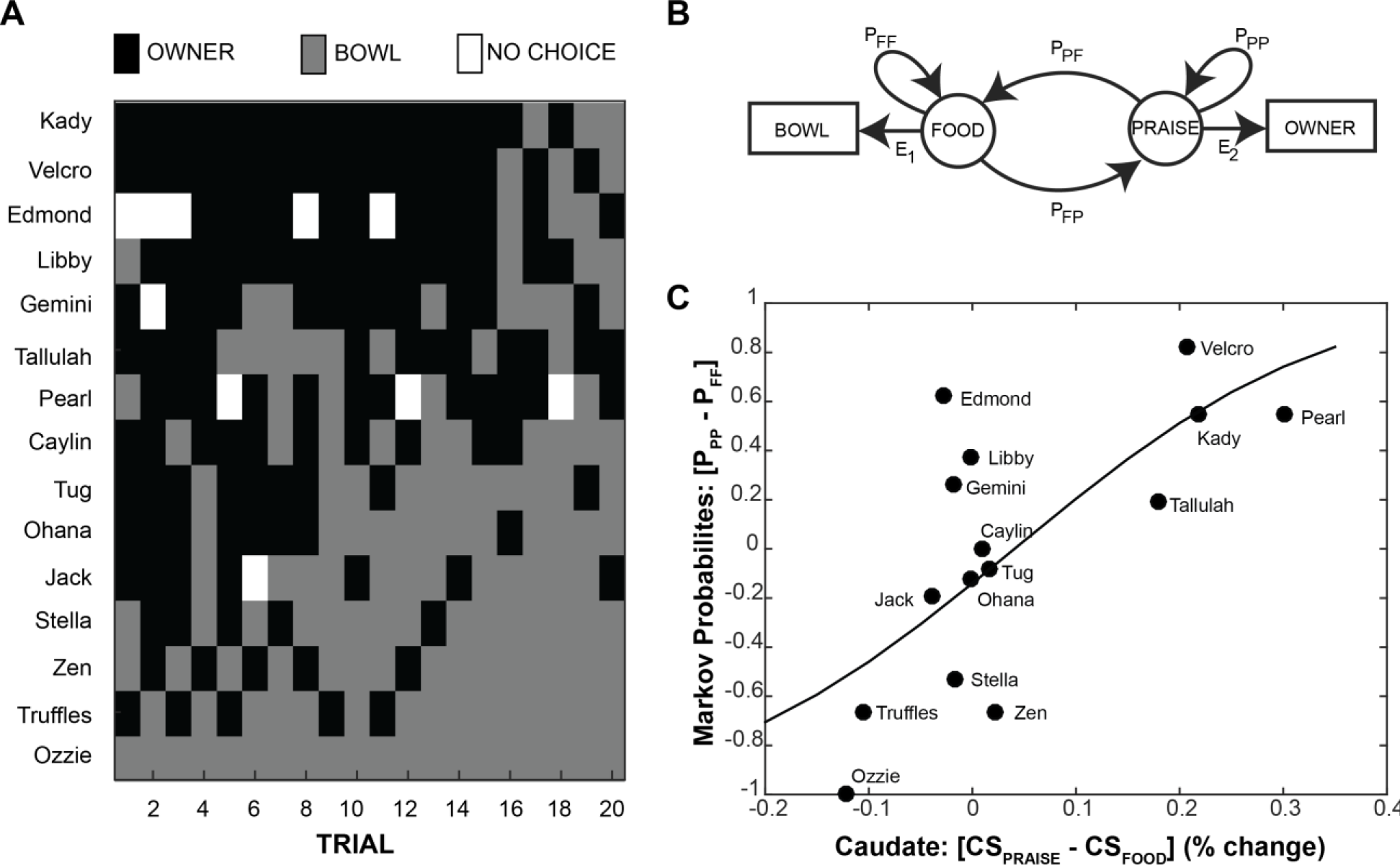
Hidden Markov model and logistic regression of neural versus behavioral preference. (*A*)Y maze choice sequences for each participant on each of the 20 test trials. (B) A schematic representation of the Hidden Markov Model used to compute transition probabilities between food bowl and owner in the behavioral Y-maze task (Experiment 3). “Food” and “Praise” represent the internal states associated with selection of the food bowl “Bowl” and owner “Owner” respectively. E represents emission probabilities, the likelihood of moving from an internal state to the matched behavior. P represents probabilities, for transitioning between states or staying in the current state. (*C*) A logit function was fit to the relationship between the differential caudate activation to expectation of praise versus food in Experiment 1 (x axis) and the differential probability of staying with owner versus food in the Markov Model (y axis). To fit the logit model, we first transformed the difference in transition probabilities into an equivalent number of owner/bowl choices, according to: N(PPP-PFF+1)/2, where N was the total number of trials in which a dog made a choice. The logit model was then fit in R using a probit link function. (Z(13)=6.24, P < 0.001, null deviance = 93.08, residual deviance = 50.10, chi square goodness of fit P < 0.001).

## DISCUSSION

The primary goal of this study was to test the hypothesis that the dog’s ventral caudate activation could serve as a stable neural predictor of individual differences in dynamic choice for food and praise. We examined neural activation in an *a priori* ventral caudate mask in 15 subjects across two imaging and one behavioral experiments. We found that individual dogs’ caudate activation to cues predicting food or praise was stable across two imaging experiments, and the caudate activation was correlated with behavior in an out-of-scanner binary choice for food bowl or owner. Given the dramatically different contexts of the MRI and the choice tasks, the predictive value of the caudate activation is striking. Based on these findings, we suggest that there is consistent neurobiological orientation toward social and food reward within individual dogs, but the degree of preference may be highly variable between individuals. Whether and how breed, rearing, and genetic profile might influence this apparently stable neurobehavioral preference are questions for further study. The answers to these questions may further illuminate the genetic history of domestic dogs, and better delineate plasticity in the neurobiology of social reward – perhaps as a model for development of social reward in humans.

While 15 is a relatively low sample size in comparison to many human imaging experiments, it compares favorably with most non-human imaging work using MRI or electrophysiology, and is consistent with prior canine fMRI studies. Importantly, we focused our analyses on repeated measures of inter-individual variability, which are less susceptible to low sample size than typical group mean analyses.

Examining first the stability of the neural activation, we found a strong correlation between the differential response to the two CSs in Experiment 1 and the response to receipt of praise vs. withholding of praise in Experiment 1. In the first experiment, there was a wide range in the differential response to CS_praise_ and CS_food_. As seen in Figs. 2 & 4, dogs clustered into three groups based on the direction of this differential. The majority of the dogs had roughly equal response to CS_praise_ and CS_food_, with a differential close to zero (i.e., within one s.d. of the sample mean). Four dogs had a grossly positive differential (i.e. praise-loving), and two dogs had a grossly negative differential (i.e. food-loving). It is obvious that dogs like food, so we assume that the differential activation to cues predicting food and praise is driven by the relative value of praise given the presence of food (cf., Cromwell, Hassani & Schultz, 2005). These results are consistent with accounts of ventral striatal function in both humans and other animals, namely, that ventral caudate encodes a signal of relative expected reward value (Knutson et al., 2000; Koeneke et al., 2008; Howe et al., 2013). Even so, this relative difference could be subject to a variety of factors like satiation, time of day, personality, and effusiveness of the owner. Importantly, while we did not find a group difference in mean response to praise and food, we did not expect one. Indeed, the high degree of individual variability in caudate response, and its ability to predict behavioral response, is our primary finding.

Experiment 2 addressed the stability of the caudate differences by repeating Experiment 1, except that the expected praise was withheld on a subset of trials. This allowed us to measure the caudate response to praise itself, given that an expectation was established by the presentation of the CS. Because the praise-withholding trials were also preceded by the same CS, the expectation was the same. We found a strong correlation between [CS_praise_ - CS_food_] in Experiment 1 and [Praise – No Praise] in Experiment 2 (Fig. 2). This shows that dogs who had a stronger caudate response to the praise-predicting cue in Experiment 1 also had a strong response to praise in Experiment 2. In other words, dogs who showed a decisive neural difference remained true to form across two different experiments. Dogs who had roughly equal responses to praise-and food-predicting cues in Experiment 1, had a range of responses to praise in Experiment 2, suggesting that they were indifferent (in the economic sense) to food and praise in these contexts. Our primary experimental interest in Experiment 2 was its use for validating the caudate signal from Experiment 1 using a different paradigm (violation versus relative predictive value) and a different temporal component of reward processing (receipt versus prediction). We did not compare caudate activation during presentation of praise-predicting and food-predicting signals in Experiment 2. These would be unlikely to strongly correlate with the same conditions from Experiment 1, given potential effects of violation trials on processing during the praise-predicting stimuli, and the potential effects of less frequent food trials (which were necessitated by study design). Indifference to two rewards could manifest by responses being more susceptible to exigencies of the paradigm and being less consistent in a dynamic choice environment, as in Experiment 3.

If the ventral caudate can be taken as a measure of choiceless utility, dogs with a strong differential should show more consistent behavior on a choice task, and that is indeed what we observed in Experiment 3 (Fig. 4). Dogs who showed a strong neural preference for food or praise showed a high differential in transition probabilities from food and owner in the behavioral task. Dogs showing higher caudate activation for CS_praise_ were less likely to switch away from the owner, and dogs showing a higher caudate activation for CS_food_ were less likely to switch away from the food bowl. Dogs who did not have a strong caudate differential were much less predictable in the behavioral task, showing a wide range of response biases.

Together, results from these three experiments suggest that a strong, stable preference for social interaction or food in dogs is supported by a consistent differential reward-related response in the ventral striatum. This differential neural response may act to constrain dynamic choice across variable contexts. Dogs without a strong differential neural response may be behaviorally ambivalent, and thus much more susceptible to a range of dynamic and sometimes unpredictable factors such as satiety, attentional shifts, environment, and owner behavior. Put simply, dogs who find food more rewarding (as measured by the neural response) will reliably choose food, dogs who find social interaction more rewarding will reliably choose social interaction, and dogs who do not find one type of stimulus more rewarding than the other will show high behavioral variability and susceptibility to situational exigencies.

Although we believe these findings suggest a general and reliable neurobehavioral preference for food and social interaction in dogs, it is important to note that different types of food and social interaction might have quite different reward values, and that these values might be relative to context. Primates devalue some food rewards in comparison to others (Brosnan & De Waal, 2003), and there is some behavioral evidence that dogs prefer physical contact with humans to verbal praise (Feuerbacher & Wynne, 2015). We emphasize the consistency of our findings, despite using different food and social reward types in the imaging (hot dog and verbal praise) and behavioral paradigms (Pupperoni and brief petting). In addition, because the reward value of food to dogs is more firmly established than the reward value of human interaction, we used high value food rewards, and relatively limited and muted social interactions. Despite this, the relative response to social versus food reward was quite high in a number of dogs. Future studies examining a wider range of rewards could be of significant interest.

On a practical level, our results emphasize the importance of social reinforcement to dogs. Thirteen of fifteen participants showed roughly equal or greater caudate activation to expectation of praise than expectation of food reward. The stability of this neural marker of preference, and its prediction of dynamic choice behavior, suggest that the majority of our participants found social interaction at least as rewarding as food. These findings are consistent with social attachment accounts of the dog-human bond (Topal et al., 1998; Palmer & Custance, 2008; Nagasawa et al., 2015) and could provide a proximal neural mechanism supporting that attachment: reward-related activity in the ventral striatum. Of note, prior association of owner with food could drive caudate response to praise, and evidence suggests that even just viewing an unfamiliar human face may activate ventral striatum in dogs (Cuaya et al., 2016). Clearly, there are multiple avenues by which humans may come to be rewarding to dogs. The strength of our current findings is in the stable neurobehavioral preference for owner over high value food in a subset of participants. Further studies might examine whether attachment history, training history, breed, or genetic profile best predict this preference.

Notably, due to the behavioral variability of dogs without a strong differential neural activation to food and praise, the Y-maze test alone could not have distinguished these dogs from those showing consistent neurobehavioral preference for food or praise. In keeping with models of choiceless utility, the differential activity in ventral striatum may be a “cleaner” signal of stable preference than can be obtained from an individual behavioral task. This suggests that neural preference tests may be useful for identifying dogs well suited to particular service roles. A dog with high preference for social reward might be best suited for certain therapeutic or assistance jobs, while a dog with less of a neural preference for social reward might be better suited for tasks that require more independence from humans, like search-and-rescue dogs or hearing-assistance dogs. Dogs without a strong neural differential could be less predictable, and thus ill-suited for specific working roles. More generally, our findings support the use of social praise as a reward in dog training (cf., McIntire & Colley, 1967; Feuerbacher & Wynne, 2012). For most dogs, social reinforcement is at least as effective as food – and probably healthier too.

## AUTHOR CONTRIBUTIONS

Conceptualization: P.F.C, A.P., M.S., and G.B. Methodology: P.F.C, A.P., M.S., and G.B. Formal Analysis: P.F.C., A.P., and G.B. Investigation: P.F.C, A.P., M.S., and G.B. Writing – Original Draft: P.F.C. and G.B. Writing – Review & Editing: P.F.C, A.P., M.S., and G.B. Visualization: P.F.C. and G.B. Funding Acquisition: G.B.

## ACKNOWLEDGMENTS

This study was performed in strict accordance with the recommendations in the Guide for the Care and Use of Laboratory Animals of the National Institutes of Health. The study was approved by the Emory University IACUC (Protocol #DAR-2001274-120814BA), and all dogs’ owners gave written consent for participation in the study. This work was supported by the Office of Naval Research (N00014-13-1-0253). G.B. and M.S. own equity in Dog Star Technologies and developed technology used in some of the research described in this paper. The terms of this arrangement have been reviewed and approved by Emory University in accordance with its conflict of interest policies. P.F.C. and A.P. claim no conflicts of interest.

## Movie 1

Participant Kady is presented with the three experimental stimuli in the mock scanner during conditioning.

## Movie 2

Participant Ohana chooses food in the Y maze task.

## Movie 3

Participant Ohana chooses owner in the Y maze task.

